# Golgi-localized GALT7 and GALT8 participate in cellulose biosynthesis

**DOI:** 10.1101/2021.04.21.440809

**Authors:** Pieter Nibbering, Romain Castilleux, Gunnar Wingsle, Totte Niittylä

**Affiliations:** Department of Forest Genetics and Plant Physiology, Umeå Plant Science Centre, Swedish University of Agricultural Sciences, 901 83 UMEÅ

**Author notes:** corresponding author: Totte Niittylä. The author responsible for distribution of materials integral to the findings presented in this article in accordance with the policy described in the Instructions for Authors (www.plantcell.org) is: Totte Niittylä.

## Abstract

Arabinogalactan protein (AGP) glycan biosynthesis in the Golgi apparatus contributes to plant cell wall assembly, but the mechanisms underlying this process are largely unknown. Here, we show that two putative galactosyltransferases -named GALT7 and GALT8 -from the glycosyltransferase family 31 (GT31) of *Arabidopsis thaliana* participate in cellulose biosynthesis. *galt7galt8* mutants show primary cell wall defects manifesting as impaired growth and cell expansion in seedlings and etiolated hypocotyls, along with secondary cell wall defects, apparent as collapsed xylem vessels and reduced xylem wall thickness in the inflorescence stem. These phenotypes were associated with a ∼30% reduction in cellulose content, a ∼50% reduction in secondary cell wall CELLULOSE SYNTHASE (CESA) protein levels and reduced cellulose biosynthesis rate. *CESA* transcript levels were not significantly altered in *galt7galt8* mutants, suggesting that the reduction in CESA levels was caused by a post-transcriptional mechanism. We provide evidence that both GALT7 and GALT8 localise to the Golgi apparatus, while quantitative proteomics experiments revealed reduced levels of the entire FLA subgroup B in the *galt7galt8* mutants. This leads us to hypothesize that a defect in FLA subgroup B glycan biosynthesis reduces cellulose biosynthesis rate in *galt7galt8* mutants.

## Introduction

Plant cell walls are necessary for plant growth and development, as well as one of the most bountiful sustainable raw materials. The main structural component in plant cell walls is cellulose, which consists of crystalline β-1,4-glucan chain microfibrils. Plant cell expansion driven growth and the final cell dimensions are dictated by turgor-driven anisotropic extension of the primary cell wall, which largely depends on cellulose microfibril (CMF) orientation and CMF interactions with cell wall matrix (Cosgrove, 2005). CMFs also contribute to the mechanical strength of secondary cell walls, which are synthesized in specific cell types, such as xylem vessels and fibers, after cell expansion (Kumar et al., 2016). Mutations affecting the biosynthesis of primary cell wall cellulose typically manifest as decreased cell expansion and growth defects in seedlings (Baskin et al., 1992; Persson et al., 2007), while a decrease in secondary cell wall cellulose levels can stunt growth, weaken stems, as well as result in thinner fiber cell walls and collapsed xylem vessels (Turner and Somerville, 1997; Taylor et al., 2003).

The β-1,4-glucan chains of CMFs are synthesized at the plasma membrane by cellulose synthases (CESAs). In *Arabidopsis thaliana* (Arabidopsis), primary cell wall cellulose is synthesised by CESA1-, CESA3- and CESA6-related proteins, while CESA4, CESA7 and CESA8 synthesise cellulose for the secondary cell walls (Taylor et al., 2003; Desprez et al., 2007; Persson et al., 2007). CESAs assemble within the catalytic core of the cellulose synthase complex (CSC) in the Golgi apparatus, from where they are transported to the plasma membrane (Crowell et al., 2009). Besides CESAs, several other proteins are involved in the *in vivo* biosynthesis of cellulose by supporting the assembly (Vain et al., 2014; Zhang et al., 2016) and trafficking of CSCs along cortical microtubules (Gu et al., 2010; Bringmann et al., 2012). A comprehensive, up-to-date account of CSC-associated proteins can be found in recent reviews by Polko and Kieber (2019) and Lampugnani et al. (2019). There is also experimental evidence implicating a role for protein glycosylation, and hence, glycoproteins, in cellulose biosynthesis.

Protein glycans can be classified as either asparagine-linked *N-*glycans or serine-, threonine- or hydroxyproline-linked *O*-glycans (Strasser, 2016). Both *N*- and *O-*glycosylation have been linked with cellulose biosynthesis. Notably, *cyt1* mutants lack the mannose-1-phosphate guanylyltransferase required for *N*-glycosylation and, subsequently, contain less cellulose than wild type (WT) embryos (Lukowitz et al., 2001). Moreover, mutant Arabidopsis embryos (*knf* and *rsw3*) containing defective alpha-glycosidases, which catalyse reactions required in *N*-linked glycan trimming, contain significantly less cellulose than WT embryos, while other cell wall polymers are only moderately affected (Burn et al., 2002; Gillmor et al., 2002). The mechanism underlying the observed cellulose defects in *cyt1, knf* and *rsw3* may be indirect, since *N*-glycans have numerous and diverse functions, including protein folding and targeting, endocytosis, signalling and modulation of protein interactions (Varki, 2017). However, a direct effect on the cellulose biosynthesis machinery is also possible. Of the known cellulose biosynthesis associated proteins, the membrane-bound endo-1,4-β-glucanase KORRIGAN1 (KOR1) was shown to be *N*-glycosylated, while the Fasciclin-like arabinogalactan 4 (FLA4) protein is both *N*- and *O-*glycosylated (Rips et al., 2014; Xue et al., 2017). KOR1 is involved in cellulose biosynthesis and the intracellular trafficking of CSCs (Nicol et al., 1998; Vain et al., 2014). Mutations in the predicted *N*-glycan sites of KOR1 showed that glycosylation affects recombinant KOR1 glucanase activity *in vitro* (Liebminger et al., 2013; Schoberer et al., 2013), and the subcellular localisation of fluorescent protein-tagged KOR1 *in vivo* (Rips et al., 2014). Glycosylation was also observed to influence the cellular trafficking of fluorescent protein-tagged FLA4, with *N*-glycosylation affecting protein exit from the endoplasmic reticulum (ER) and *O-*glycosylation influencing plasma membrane localisation (Xue et al., 2017). FLA4 is a member of the arabinogalactan protein (AGP) family, which includes hydroxyproline-rich glycoproteins that are abundant in the plasma membrane and cell wall (Showalter et al., 2010).

AGPs in plants are subdivided into the classical AGPs, lysine-rich AGPs, AG peptides, Fasciclin-like AGPs (FLAs) and other chimeric AGPs (Showalter et al., 2010). Arabidopsis contains 21 FLAs, which are further classified into four subgroups (labelled A, B, C and D) based on their amino acid sequence and domain structures (Johnson et al., 2003). Of the AGPs characterized until now, only members of the FLA family have been linked to cellulose biosynthesis. For example, *fla4* (also known as *salt overly sensitive 5*) mutants exhibit root cell expansion defects when grown on media with a high NaCl content (Shi et al., 2003). The cell expansion phenotype was associated with conditional cellulose biosynthesis defects in primary cell walls; more specifically, *fla4* seedlings incorporated less ^14^C-glucose into cellulose under NaCl stress (Basu et al., 2016). FLAs have also been linked to secondary cell wall cellulose biosynthesis, with MacMillan et al. (2010) demonstrating that the simultaneous mutation of *FLA11* and *FLA12* results in reduced cellulose content in the inflorescence stem of Arabidopsis. It is not known how FLAs influence cellulose biosynthesis, but structural roles in cell wall matrix assembly, direct interaction with CSCs and receptor-mediated signalling mechanisms have been proposed (Tan et al., 2013; Xue et al., 2017; Seifert, 2018). In general, it has been challenging to establish mechanistic relationships between AGP functionality and plant cell wall assembly. The glycans attached to the AGP protein core are thought to be important for functionality, and the characterization of the enzymes responsible for glycan biosynthesis may provide insight into AGP function and their functional redundancy (Knoch et al., 2014).

AGP glycans consist of a β-1,3-galactan backbone with the β-1,6-linked side chains including galactose, arabinose, glucuronic acid, rhamnose and fucose (Tan et al., 2010; Tryfona et al., 2010; Tryfona et al., 2012; Kitazawa et al., 2013). The *O*-glycosylation of AGPs starts in the ER, where proline hydroxylation by proly-4-hydroxylases occurs, after which the first galactose may be attached to hydroxyproline (Hieta and Myllyharju, 2002; Rose and Lee, 2010; Hijazi et al., 2014). After exiting the ER, glycosylation continues in the Golgi apparatus, where a group of glycosyl transferases (GTs) synthesize arabinogalactan. One of the key GT families involved in AGP glycosylation is the glycosyl transferase 31 (GT31) family, which includes enzymes that participate in the synthesis of the β-1,3-galactan backbone. The Arabidopsis GT31 family consists of 33 members, of which 20 were initially suggested to be involved in the glycosylation of AGPs based on the presence of a putative β1,3-galactosyltransferase domain (Qu et al., 2008). Phylogenetic analyses have since subdivided these twenty GT31s into six clades, with the assignments based on biochemical activity predictions from sequence similarity to GT31s characterized from *Homo sapiens*. Clades I-II were predicted to have β1,3-galactosyl transferase activity, clade III was predicted to have β1,3-*N*-acetylglucosaminyltransferase activity, clade IV was predicted to have β1,3-galactosyl transferase activity for *N*-glycosylation, and clades V-VI were predicted to have hydroxyproline *O*-galactosyltransferase activity *(Qu et al*., *2008)*. Some of these predictions have been experimentally validated; for example, GALACTOSYLTRANSFERASE1 (GALT1) from clade V was found to be a β–1,3-galactosyltransferase essential for the attachment of β1,3-galactose residues to *N*-glycans (Strasser et al., 2007). Moreover, GALT2-GALT6 (clade V-VI) and hydroxyproline O-galactosyltransferases (HPGTs, Clade III) have been shown to be hydroxyproline O-galactosyltransferases (Basu et al., 2013; Basu et al., 2015a; Basu et al., 2015b; Ogawa-Ohnishi and Matsubayashi, 2015). As such, *galt2 – galt6* mutants displayed multiple growth defects, including reduced root hair growth, reduced seed coat mucilage and seed set, along with accelerated leaf senescence and reduced root growth under salt stress (Basu et al., 2015a; Basu et al., 2015b). Furthermore, triple *hpgt* mutants (*hpgt1,2,3*) with only 10% of wild-type HPGT activity remaining exhibited reduced overall growth rate, but had longer lateral roots and root hairs, as well as increased radial expansion of cells in the root tip (Ogawa-Ohnishi and Matsubayashi, 2015). The *Arabidopsis thaliana* GT31 enzymes from Clade IV have not been characterized, but cotton orthologs suggest that Clade IV possesses β–1,3-galactosyltransferase activity (Qin et al., 2017). The enzyme GALT31A (Clade II) was initially characterized to have β–1,6-galactosyltransferase activity (Geshi et al., 2013), but a recent study indicated that it is actually a β–1,3-galactosyltransferase (Ruprecht et al., 2020). Mutations in Arabidopsis *GALT31A* were found to arrest embryonic development at the globular stage, which indicates that this protein and β– 1,3-galactan linkages it catalyses are crucial in plants (Geshi et al., 2013). A close Clade II homolog of GT31A called KAONASHI4 (KNS4) was also shown to be a β–1,3-galactosyltransferase, with *kns4* mutants displaying reduced fertility, shorter seed pods and reduced seed set due to defects in pollen exine development (Suzuki et al., 2017). Thus, the biosynthetic activity has been established for eleven out of the twenty putative AGP glycosylating GT31s. Moreover, the characterization of corresponding Arabidopsis mutants has established that GT31s function in a wide variety of growth and developmental processes in plants. However, the *in vivo* targets of these enzymes have not yet been established; therefore, the link between GT31 activity and plant phenotype often remains unclear. Here we present results on two previously uncharacterized GT31 family enzymes from clade I, and describe their involvement in primary and secondary cell wall cellulose biosynthesis, possibly through β–1,3-galactosylation of the FLA subgroup B.

## Results

### *GALT7* and *GALT8* mutants display primary and secondary cell wall defects

The analysis of co-expressed genes has proven effective for identifying functionally related cell wall biosynthesis genes in Arabidopsis (Aoki et al., 2007; Mutwil et al., 2009; Ruprecht et al., 2011). In an effort to investigate the role of β1,3-galactans in plant cell walls, we recently identified two Golgi-localized exo-β1,3-galactosidases involved in cell expansion and root growth in Arabidopsis (Nibbering et al., 2020). In order to identify enzymes participating in β1,3-galactosidase substrate synthesis *in vivo*, we searched for co-expressed β1,3-galactosyltransferase genes using the publicly available Arabidopsis co-expression tools (https://atted.jp/ and https://genevestigator.com/). This resulted in the identification of two co-expressed candidates from the GT31 family, namely, AT1G05170 and its close sequence homolog AT4G26940. To follow the published *GT31* family nomenclature we named these genes *GALT7* and *GALT8*, respectively. Both *GALT7* and *GALT8* are expressed throughout the Arabidopsis plant, with the bottom internodes of the inflorescence stem showing the highest expression (http://bar.utoronto.ca/eplant/).

To investigate the function of *GALT7* and *GALT8*, we obtained T-DNA insertion mutants for both genes (Fig. 1A). Homozygous single *GALT7* or *GALT8* T-DNA insertion lines did not show obvious phenotypical differences relative to WT plants when grown under controlled conditions, but double *galt7galt8* mutants demonstrated clear growth defects. Allelic *galt7-1/galt8-1* and *galt7-2/galt8-2* seedlings both exhibited shortened primary roots and etiolated hypocotyls (Fig. 1B-C). Quantification of the etiolated hypocotyl length confirmed a significant reduction in both double mutant lines relative to WT seedlings (Fig. 1D). Since hypocotyl elongation is primarily driven by cell expansion rather than cell division, this observation supports that GALT7 and GALT8 are involved in primary cell wall biosynthesis and cell expansion. At maturity, the inflorescences of 10-week-old *galt7-1/galt8-1* and *galt7-2/galt8-2* mutants were clearly stunted, indicating that both GALT7 and GALT8 also function after the seedling stage (Fig. 1E). Cross sections of the 10-week-old inflorescence stems of the *galt7galt8* lines revealed reduced fiber cell wall thickness and the occasional collapse of xylem vessels (Supplementary Figs. 1 and 2). Moreover, transmission electron microscope (TEM) analysis of the cross sections confirmed thinner secondary cell walls in the vessels, vascular fibers and interfascicular fibers of the *galt7galt8* mutants relative to WT plants (Fig. 2). No difference was observed in the TEM images of *galt7* or *galt8* single mutant stem cross sections (Supplementary Fig. 3). To establish a causal link between the phenotypes and T-DNA insertions, the *galt7-1/galt8-1* mutant was transformed with either *GALT7-YFP* or *GALT8-YFP* under the control of a native promoter. Both constructs complemented the etiolated hypocotyl phenotype (Supplementary Fig. 4A/B) and the stunted growth in soil-grown plants (Supplementary Fig. 5). These results establish that GALT7 and GALT8 act redundantly during both primary and secondary cell wall biosynthesis.

**Figure 1.**
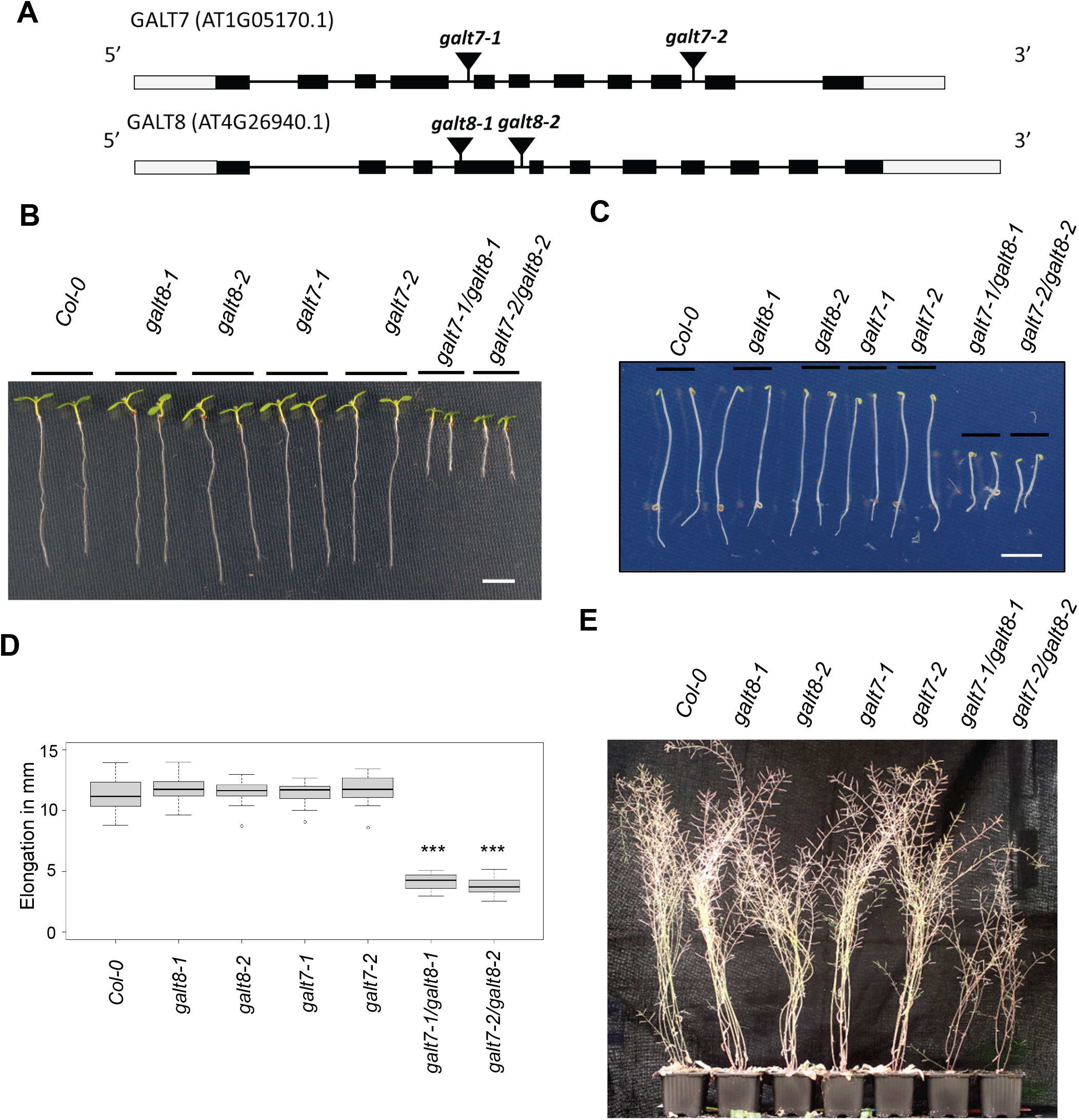
Characterisation of the *galt7* and *galt8* T-DNA insertion lines. **(A)** Schematic diagram of the *GALT7* and *GALT8* gene structure along with the T-DNA insertion sites. The grey and black boxes indicate untranslated and translated regions, respectively. **(B)** Seven-day-old *Col-0* and *galt* seedlings grown on ½ MS agar plates (scale bar = 5mm) **(C)** Four-day-old etiolated *Col-0* and *galt* seedlings grown on ½ MS agar plates in the dark (scale bar = 5mm) **(D)** Hypocotyl length of four-day-old etiolated *Col-0* and *galt* seedlings. In the boxplot, dark horizontal lines represent the median, the two grey boxes denote the 25th and 75th percentiles, the whiskers denote the 1.5 interquartile range limits, and the dots are outliers. ***P < 0.001 (unpaired t test, *n* = 25-30 biological replicates). **(E)** Ten-week-old senescing *Col-0* and *galt* lines.

**Figure 2:**
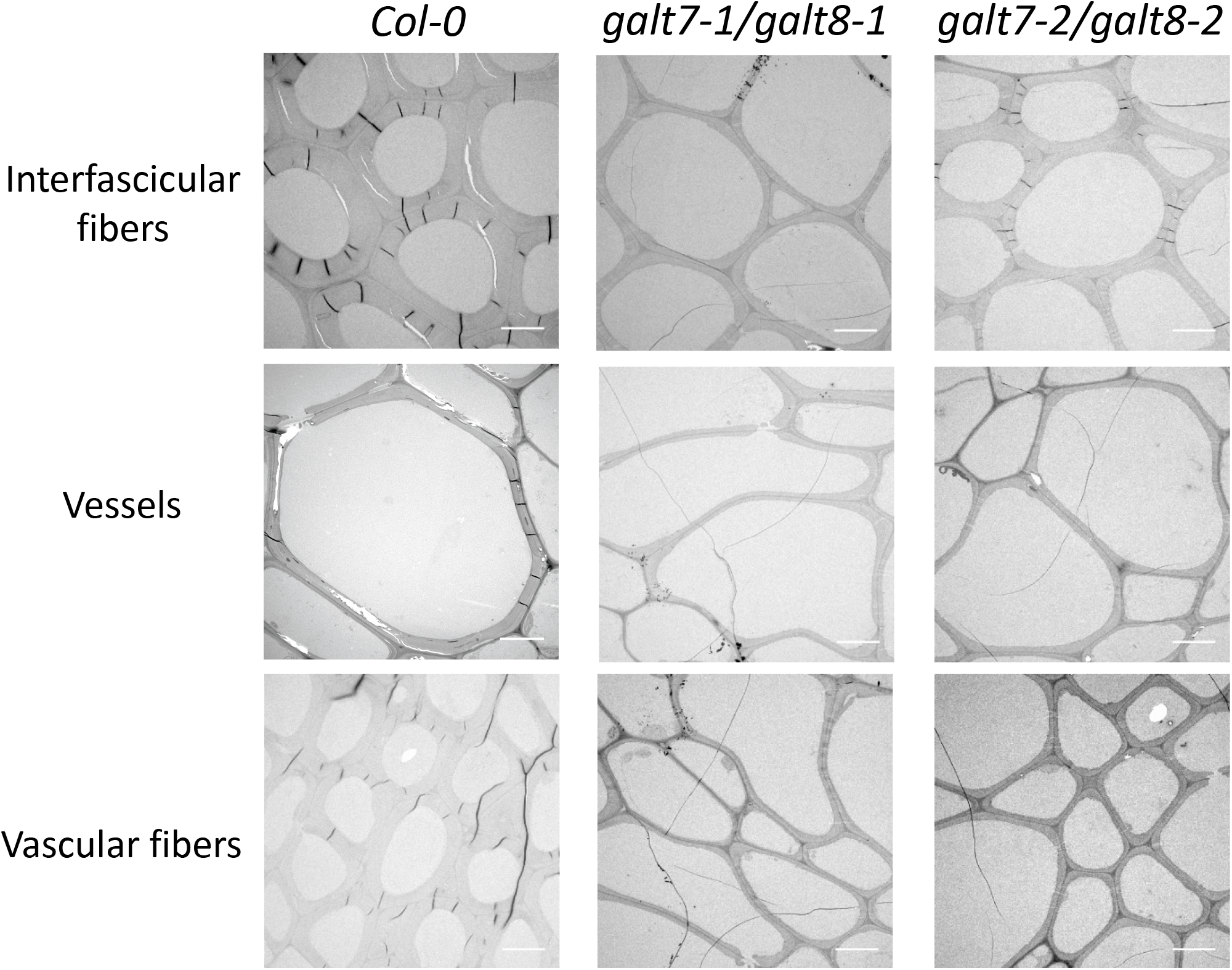
Transmission electron microscope (TEM) images of interfascicular fibers, vessels and vascular fibers from cross sections of 10-week-old inflorescence stems of *Col-0, galt7-1/galt8-1 and galt7-2/galt8-2*. Scale bar = 5 µm

### *galt7galt8* mutants demonstrate defective cellulose biosynthesis

To investigate the cause of the decreased cell expansion and wall defects, we analysed the cell wall composition of the mutants. The single *galt7* and *galt8* mutants did not differ significantly from WT lines in terms of cellulose content, whereas the *galt7galt8* lines showed a ∼30% decrease in cellulose content relative to WT lines in both 10-week-old inflorescence stems and 4-day-old dark-grown hypocotyls (Table 1, Supplementary Fig. 6). The cellulose content was fully restored in the stems of the *GALT7-YFP* or *GALT8-YFP* complemented *galt7galt8* lines, which confirms a causal link between both GALT7 and GALT8 and cellulose synthesis (Supplementary Fig. 7). Furthermore, the *galt7galt8* inflorescence stems showed higher levels of rhamnose, xylose, mannose, galacturonic acid, methylated glucuronic acid and lignin than the WT stems (Table 1). This increase in hemicellulosic sugars and lignin per dry weight is likely a consequence of the decrease in cellulose. We also noted that the amount of galactose did not differ between the mutant and WT lines, which suggests that GALT7 and GALT8 are not major contributors to the cell wall galactose pool (Table 1).

**Table 1:** Crystalline cellulose, hemicellulosic/pectic sugar and lignin content of 10-week-old stems from *Col-0* and *galt* lines (%, dry weight). Unpaired *t*-test, *P < 0.05, **P < 0.01, ***P < 0.001 (*n* = 5 biological replicates)

|  | *Col-0* | *galt8-1* | *galt8-2* | *galt7-1* | *galt7-2* | *galt7-1/galt8-1* | *galt7-2/galt8-2* |
| --- | --- | --- | --- | --- | --- | --- | --- |
| **Crystalline cellulose** | 38.2 ± 0.7 | 36.0 ± 1.4 | 40.5 ± 3.1 | 40.2 ± 0.9 | 40.1 ± 1.0 | **27.2 ± 1.2\*\*\*** | **27.8 ± 0.7\*\*\*** |
| **Arabinose** | 1.25 ± 0.03 | 1.30 ± 0.03 | 1.36 ± 0.05 | 1.25 ± 0.02 | 1.24 ± 0.02 | 1.25 ± 0.03 | 1.33 ± 0.03* |
| **Rhamnose** | 1.32 ± 0.04 | 1.42 ± 0.03 | 1.45 ± 0.08 | 1.35 ± 0.03 | 1.33 ± 0.02 | 1.46 ± 0.04* | 1.59 ± 0.05** |
| **Fucose** | 0.92 ± 0.02 | 0.98 ± 0.02* | 0.95 ± 0.02 | 0.94 ± 0.01 | 0.90 ± 0.02 | 0.95 ± 0.03 | 0.95 ± 0.02 |
| **Xylose** | 15.09 ± 0.54 | 14.78 ± 0.46 | 15.92 ± 1.04 | 15.48 ± 0.64 | 15.34 ± 0.40 | 17.21 ± 0.48* | 18.99 ± 0.65** |
| **Mannose** | 1.83 ± 0.08 | 1.90 ± 0.05 | 1.89 ± 0.12 | 1.78 ± 0.07 | 1.92 ± 0.06 | 2.06 ± 0.05* | 2.14 ± 0.07* |
| **Galactose** | 1.74 ± 0.03 | 1.78 ± 0.04 | 1.83 ± 0.06 | 1.72 ± 0.01 | 1.70 ± 0.03 | 1.77 ± 0.04 | 1.81 ± 0.04 |
| **Galacturonic acid** | 3.94 ± 0.17 | 4.38 ± 0.10 | 4.57 ± 0.28 | 3.98 ± 0.07 | 4.16 ± 0.09 | 4.84 ± 0.24* | 5.31 ± 0.20*** |
| **Glucose** | 4.76 ± 0.15 | 5.28 ± 0.36 | 5.17 ± 0.18 | 4.29 ± 0.15 | 4.71 ± 0.32 | 5.02 ± 0.32 | 4.46 ± 0.20 |
| **Glucuronic acid** | 1.27 ± 0.04 | 1.32 ± 0.02 | 1.32 ± 0.05 | 1.28 ± 0.02 | 1.30 ± 0.03 | 1.27 ± 0.03 | 1.35 ± 0.02 |
| **Methyl Glucuronic acid** | 1.68 ± 0.08 | 1.77 ± 0.04 | 1.75 ± 0.09 | 1.68 ± 0.06 | 1.75 ± 0.05 | 2.09 ± 0.05** | 2.18 ± 0.05*** |
| **Matrix polysaccharide total** | 33.80 ± 1.10 | 34.91 ± 0.89 | 36.22 ± 1.83 | 33.76 ± 0.82 | 34.36 ± 0.85 | 37.90 ± 1.07* | 40.09 ± 1.05** |
| **Lignin** | 21.85 ± 0.57 | 21.73 ± 0.40 | 21.57 ± 0.62 | 23.66 ± 0.47 | 23.06 ± 0.81 | 25.14 ± 0.72* | 25.19 ± 1.09* |

To investigate the cause of the cellulose defect observed in *galt7galt8* mutants, we analysed the expression levels of primary cell wall *CESA1, CESA3* and *CESA6* in 4-day-old dark-grown hypocotyls, along with the expression of secondary cell wall *CESA4, CESA7* and *CESA8* in the inflorescence stem, using quantitative PCR (qPCR). The WT and *galt7galt8* lines did not differ significantly in the expression of primary or secondary cell wall *CESA* transcripts, indicating that the cellulose defect is not caused by transcriptional deficiency (Fig. 3A). To investigate CESA protein levels, secondary cell wall CESA4, 7 and 8 were analysed using quantitative mass spectrometry and isotope-labelled standard peptides according to a method presented by Zhang et al. (2018). This assay revealed significant reductions (∼50%) in secondary cell wall CESA protein levels in the *galt7galt8* mutants (Fig. 3B). This conclusion was also supported by reduced band intensity of CESA4, 7 and 8 in *galt7galt8* compared to WT on Western blots probed with CESA isoform specific antibodies (Supplementary Fig. 8). Differences in the rates of primary cell wall cellulose biosynthesis were investigated by comparing the rates at which growth media-supplied ^13^C-glucose was incorporated into cellulose in *galt7galt8* and WT seedlings. A time course experiment revealed a significant reduction in the amount of ^13^C in the crystalline cellulose fraction of *galt7galt8* cell walls at all time points compared to WT, while total ^13^C uptake was significantly lower in *galt7galt8* mutants relative to the WT line only at the 16h time point (Fig. 3C and 3D). These results suggest that the reduced cellulose content observed in *galt7galt8* mutants is caused by a post-transcriptional mechanism which reduces CESA levels and, subsequently, the rate of cellulose biosynthesis.

**Figure 3:**
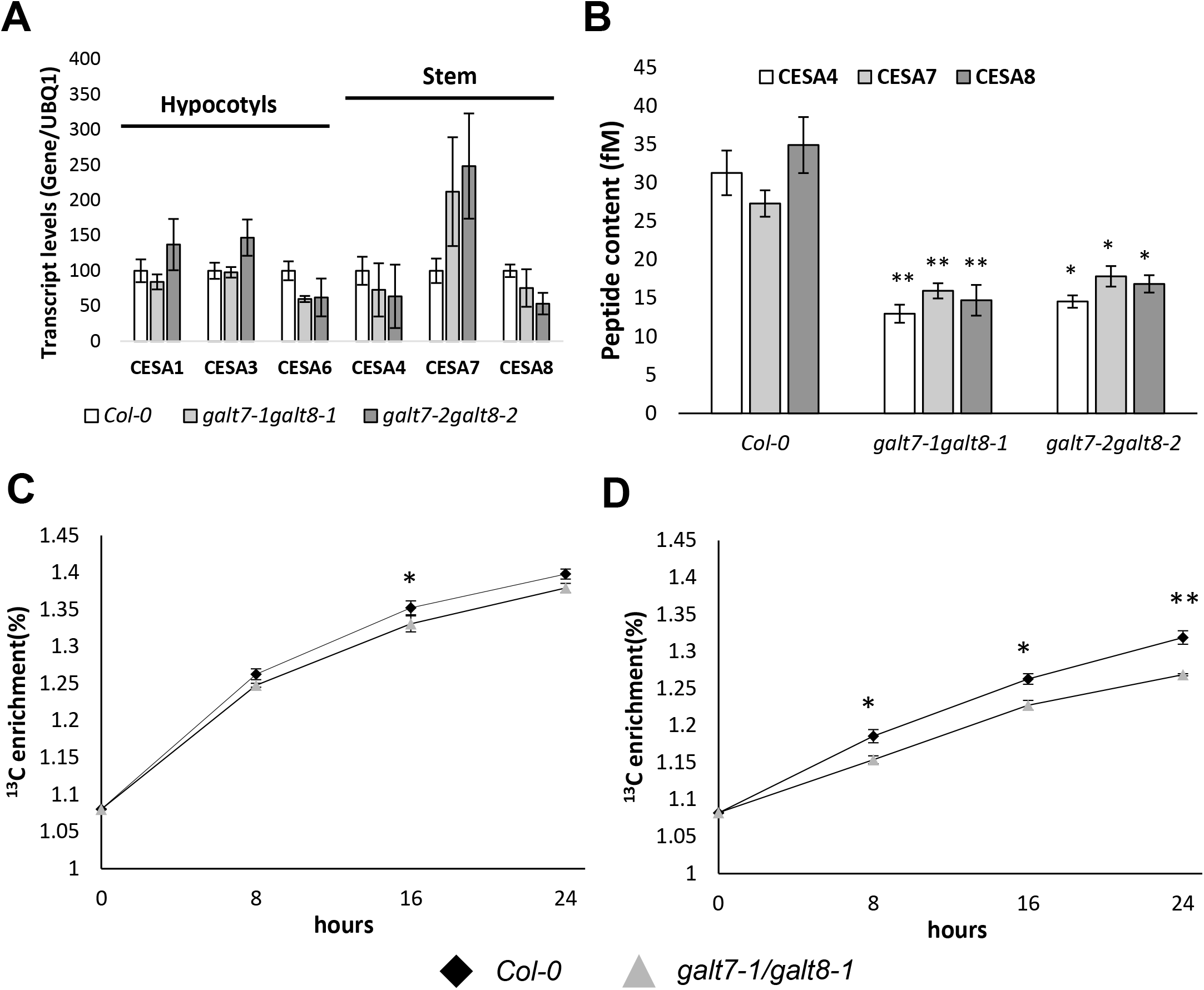
*CESA* transcript, CESA protein levels, and cellulose biosynthesis rates measured in *Col-0* and *galt7galt8* plants. **(A)** Expression levels of *CESA1, 3* and *6* in 4-day-old etiolated seedlings and *CESA4, 7* and *8* in the inflorescence stems analyzed by RT-qPCR (Mean ± SE, *n*= 3 biological replicates). **(B)** Secondary cell wall CESA peptide levels from the first 10 cm of 20 cm long inflorescence stems from *Col-0* and the *galt7galt8* mutants. Significance is calculated with a two sided student’s Unpaired t-test. *P-value = * ≤0.05, **P ≤0.01. (Mean ± SE, *n*= 4 biological replicates). **(C)** Total ^13^C enrichment in eight-day old Col-0 and *galt7galt8* seedlings after ^13^C-glucose supply. **(D)** ^13^C enrichment in crystalline cellulose in eight-day old Col-0 and *galt7galt8* seedlings. ^13^C enrichment was determined with an Elemental Analyzer-Isotope Ratio Mass Spectrometer (EA-IRMS) and is shown as a percentage of total carbon. The data represents 3 biological replicates (Mean ± SD) of ∼1000 pooled seedlings per time point. Significance is calculated with a two sided student’s Unpaired *t*-test. *P-value = * ≤0.05, **P ≤0.01.

### GALT7-YFP and GALT8-YFP are Golgi-localized

To provide an explanation for the reduced secondary cell wall CESA levels and reduced cellulose biosynthesis rates observed in the *galt7galt8* seedlings, we first established the subcellular localisation of GALT7 and GALT8 using the *GALT7-YFP* and *GALT8-YFP* lines. In the epidermal cells of the root elongation zone of four-day old seedlings, both GALT7-YFP and GALT8-YFP appeared to be localised to the Golgi apparatus (Fig. 4). To validate the Golgi localisation, the lines were crossed with the *cis*-Golgi marker SYP32-mCherry and the Trans Golgi Network (TGN) marker SYP43-mCherry (Uemura et al., 2004; Geldner et al., 2009). Both GALT7-YFP and GALT8-YFP co-localised with the Golgi markers, and showed stronger overlap with SYP32-mCherry than with SYP43-mCherry (Fig. 4A). It was difficult to quantify the extent of co-localization due to the weakness of the YFP signal. Therefore, to confirm that the GALT7/8-YFP proteins were localised to the Golgi rather than the TGN, the lines were treated with brefeldin A for 30 minutes. Brefeldin A causes the fusion of early Golgi cisternae and the endoplasmic reticulum, while Golgi cisternae facing the trans side of the Golgi and TGN are dispersed to the cytoplasm (Nebenfuhr et al., 2002). After the Brefeldin-A treatment, both GALT7-YFP and GALT8-YFP co-localised with SYP32-mCherry, which confirms a *cis-*Golgi and/or medial-Golgi localisation (Fig. 4B).

**Figure 4:**
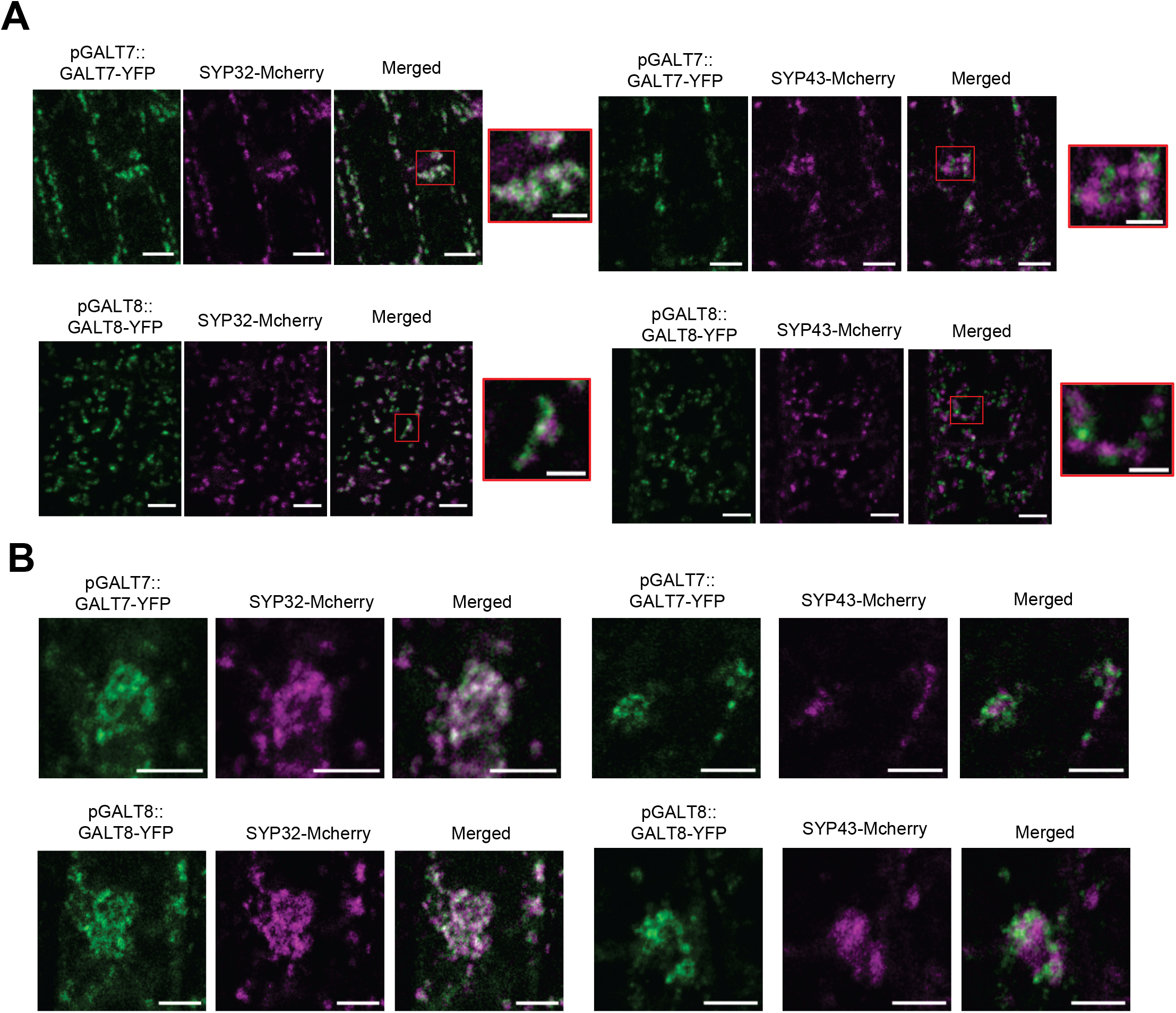
Co-localization of GALT7-YFP and GALT8-YFP with mCherry Golgi markers. **(A)** Confocal laser scanning microscope images of root epidermis cells in Arabidopsis seedlings stably expressing pGALT7:GALT7-YFP or pGALT8:GALT8-YFP and the cis-Golgi marker SYP32-mCherry or the Trans Golgi Network (TGN) marker SYP43-mCherry. Images were captured with a Zeiss LSM880 confocal microscope. Scale bars = 5µm. (2µm for zoomed image) **(B)** Confocal laser scanning microscope images of root epidermis cells in Arabidopsis seedlings stably expressing pGALT7:GALT7-YFP or pGALT8:GALT8-YFP and the cis-Golgi marker SYP32-mCherry or the Trans Golgi Network (TGN) marker SYP43-mCherry after treatment with 25µM Brefeldin A for 30 min. Images were captured with a Zeiss LSM880 confocal microscope. Scale bars = 5µm

### *galt7galt8* mutants show decreased levels of AGP-linked β-1,3 galactan

In plants, β1,3-galactan is primarily associated with AGPs, and AGP glycosylation takes place in the Golgi. For this reason, we measured the amounts of β1,3-galactan in *galt7galt8* and WT lines using β-Yariv phenylglycoside. β-Yariv binds specifically to β1,3-galactan consisting of a minimum of seven monomeric units, and can be used to quantify AGP galactans in plant extracts (Kitazawa et al., 2013). β-Yariv precipitation, followed by spectrophotometric quantification, revealed that *galt7galt8* lines contained ∼35%, ∼22%, ∼25% less AGPs in the inflorescence stem, 7-day-old seedlings, and 4-day-old etiolated hypocotyls, respectively, relative to the corresponding WT plants (Fig. 5A). The inflorescence stems were used for further AGP characterization. Soluble AGPs were extracted from the stems with hot water and analysed using a β-Yariv gel diffusion assay and gel electrophoresis. The gel diffusion assay confirmed a reduction in β-Yariv accessible β-1,3 galactan from *galt7galt8* relative to WT levels (Fig. 5B). The agarose gel electrophoresis revealed no major changes in the MW distribution of AGPs detected from *galt7galt8* mutant and WT stems leading us to hypothesise that GALT7 and GALT8 act on a subset of AGPs (Fig. 5C).

**Figure 5.**
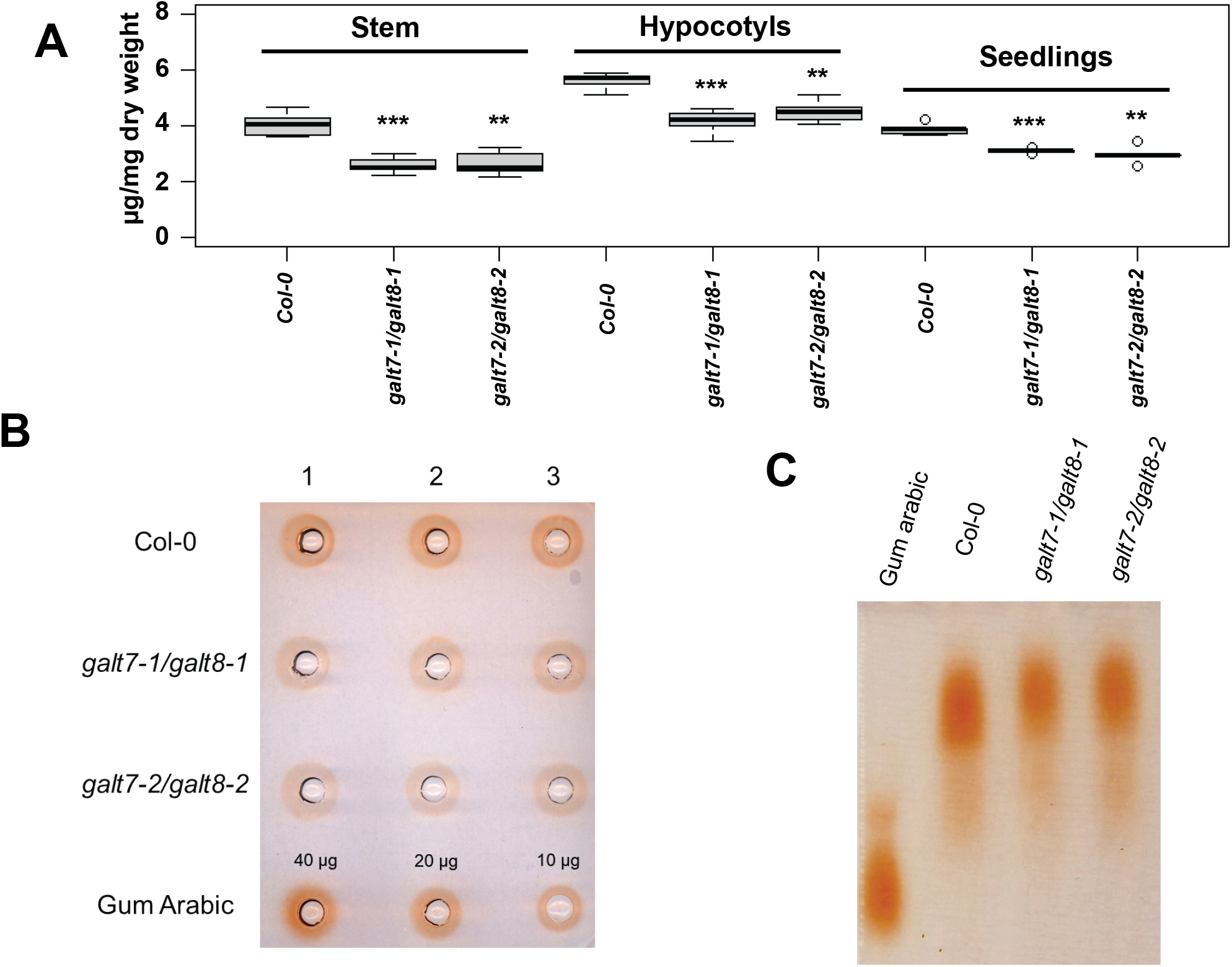
β-Yariv quantification and characterization of AGPs from Col-0 and *galt7galt8* lines. **(A)** β-Yariv precipitated AGP quantification from the bottom 10 cm of inflorescence stem, 4-day old etiolated hypocotyls and 7-day old seedlings of Col-0 and *galt7galt8* lines. β-Yariv binding to Gum Arabic was used as standard. The boxplot’s dark horizontal lines represent the median, the two grey boxes the 25th and 75th percentile, the whiskers the 1.5 interquartile limits and the dots the outliers. **P < 0.01, ***P < 0.001 (unpaired t test, *n* = 5 biological replicate). **(B)** β-Yariv gel diffusion assay of water soluble AGPs from Col-0 and *galt7galt8* lines extracted from the first 10 cm of inflorescence stem. Numbers 1, 2 and 3 indicate biological replicates. **(C)** Agarose gel electrophoresis and β-Yariv staining of water soluble AGPs from Col-0 and *galt7galt8* lines extracted from the first 10 cm of inflorescence stem. 20 µg of Gum Arabic was loaded as a control.

To further elucidate the connection between cellulose biosynthesis, GALT7&8 and AGPs, we investigated whether defective cellulose biosynthesis alone impacts AGP levels by quantifying the β-Yariv precipitated AGPs from the inflorescence stems of the secondary cell wall *cesa4* mutant. The results revealed a ∼15% reduction in AGP levels in the *cesa4* mutants, which is a less pronounced decrease than what was observed in *galt7galt8* mutants, but points to a reciprocal relationship between AGPs and cellulose biosynthesis and/or cellulose content (Supplementary Fig. 9). Therefore, it is possible that a primary defect in cellulose biosynthesis associated AGPs in *galt7galt8* mutants may lead to a further reduction in the total AGP levels.

### *galt7galt8* mutants show reduced FLA subgroup B levels

Protein glycosylation can affect protein folding, subcellular targeting, and protein turnover (Strasser, 2016; Xue et al., 2017). Hence, we hypothesised that the lack of GALT7 and GALT8 activity may affect the levels of AGPs that are relevant to cellulose biosynthesis. To identify such changes, we designed a proteomics experiment to quantify differences in the relative peptide amounts from trypsin-digested cell wall and membrane-associated protein extracts from WT and *galt7galt8* inflorescence stems (Supplementary table S1). This analysis identified 16 and 133 proteins that showed increased and decreased levels, respectively, at the *p ≤* 0.01 significance level (two-sided Student’s T-test) in *galt7galt8* lines relative to WT lines (Supplementary table S2). Several FLAs were among the peptides with the most significant changes. Moreover, we were able to quantify 12 out of the 21 FLAs found in Arabidopsis (Fig. 6) (Schultz et al., 2002; Johnson et al., 2003). Interestingly, *galt7galt8* mutants contained significantly less peptides derived from FLA15-FLA18 than the WT lines; these proteins form their own FLA subgroup B based on domain structure, sequence similarity and phylogenetic relationship (Johnson et al., 2003). In addition, *galt7galt8* mutants demonstrated a modest increase in FLA12 relative to WT lines, while FLA1, 2, 7, 8, 10, 11 and 13 levels did not differ significantly between the *galt7galt8* and WT lines (Fig. 6). These results make the FLA subgroup B a strong candidate as a component in cellulose biosynthesis *in vivo*, and leads us to hypothesise that GALT7 and GALT8 synthesise AG glycans on these specific FLAs.

**Figure 6:**
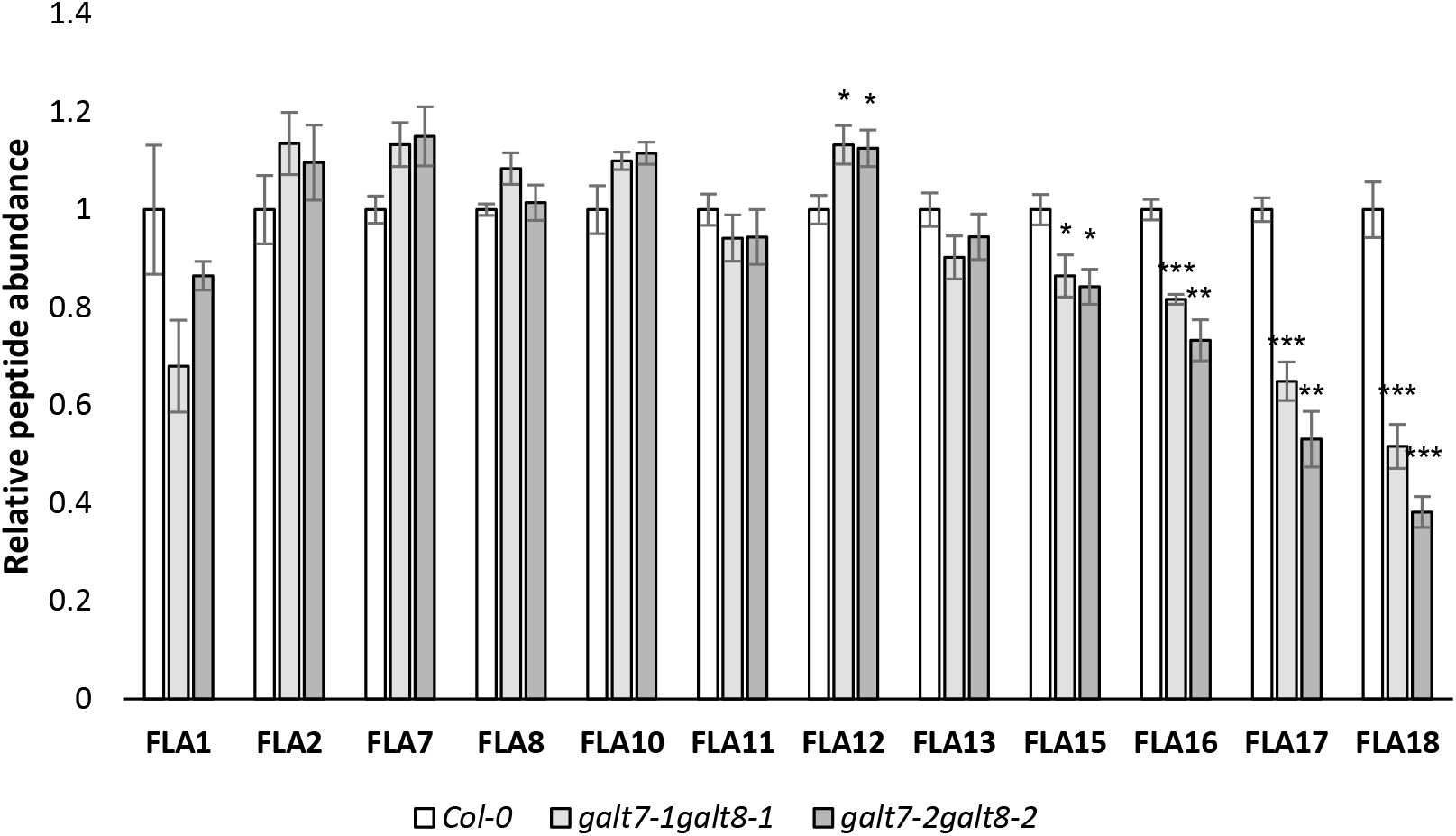
Quantification of Fasciclin-like AGPs in the stems of Col-0 and *galt7galt8* lines. The relative abundance of FLA derived peptides after trypsin digestion of cell wall and membrane associated proteins in stems of *Col-0, galt7-1galt8-1 and galt7-2galt8-2*. Peptide abundances were normalized based on total raw abundance of all the detected peptide peaks. The data represents Mean + SE (n=4 biological replicates, Two-sided student’s *t*-test, * ≤0.05, ** ≤0.01, *** ≤0.001

## Discussion

We identified two putative β1,3-galactosyl transferase genes from the GT31 family, named *GALT7* and *GALT8*, based on the co-expression of recently characterised Arabidopsis exo-β1,3-galactosidase genes (Nibbering et al., 2020). The characterization of GALT7 and GALT8 established that they are located in the *cis*-Golgi and/or medial-Golgi and act redundantly during both primary and secondary cell wall biosynthesis. *galt7galt8* mutants showed stunted seedling growth, cell expansion defects in etiolated hypocotyls and shortened inflorescence stems. When compared to WT plants, the *galt7galt8* mutants showed thinner and weaker (as evidenced by xylem vessel collapse) stem xylem fiber and vessel cell walls. The seedling and etiolated hypocotyl defects are similar to what has previously been observed in mutants with impaired primary cell wall cellulose biosynthesis (Persson et al., 2007), while the thinner xylem cell walls in the stem, along with the collapsed xylem vessels, are characteristic of secondary cell wall cellulose biosynthesis defects (Turner and Somerville, 1997; Taylor et al., 2003). These cell wall defects, when considered together with the reduced cellulose content in both *galt7galt8* seedlings and stems, indicate that GALT7 and GALT8 are involved in cellulose biosynthesis. Observations of decreased secondary cell wall CESA levels in inflorescence stems and reduced cellulose biosynthesis rates in *galt7galt8* seedlings further support this hypothesis.

Reduced β-Yariv binding in *galt7galt8* mutants and the Golgi localization of both GALT7-YFP and GALT8-YFP suggest that these proteins are involved in the synthesis of AGPs via galactosyltransferase activity. The simultaneous reduction in cellulose and AGPs observed in *galt7galt8* mutants raises the possibility that GALT7 and GALT8 act on a specific subgroup of AGPs that are associated with cellulose biosynthesis. In light of this claim, it is worth noting that even a 90% decrease in β-Yariv precipitable AGPs in the hydroxyproline galactosyltransferase mutant *hpgt1,2,3* only had a slightly negative effect on the growth of Arabidopsis seedlings or young rosette-stage plants (Ogawa-Ohnishi and Matsubayashi, 2015). In comparison, the ∼22-25% reduction in β-Yariv binding measured in *galt7galt8* seedlings resulted in strong cell expansion defects and reduced root growth. Furthermore, Zhang et al. (2021) recently characterised a quintuple hydroxyproline galactosyltransferase mutant *galt2galt3galt4galt5galt6*, which exhibited a ∼80% decrease in β-Yariv precipitable AGPs in rosette leaves and a 50-60% decrease in the stems and siliques. The quintuple mutant showed reduced overall growth, impaired root growth, abnormal pollen, shorter siliques, and reduced seed set, but the phenotype is still relatively mild in comparison to the strong growth defects observed in the *galt7galt8* mutants. This further supports the hypothesis that the involvement of GALT7 and GALT8 in cellulose biosynthesis is linked to a specific AGP glycosylation process rather than AGP glycosylation in general. Based on similar reasoning, it also seems unlikely that GALT7/8 would be involved in *N*-glycosylation, as mutants lacking GALT1 -a β1,3-galactosyl transferase essential for adding a β1,3-galactose to *N*-glycans -did not display obvious phenotypic differences when compared to WT plants (Strasser et al., 2007).

Mass spectrometry analyses of trypsin-digested cell wall and membrane-associated protein extracts from inflorescence stems revealed that all of the proteins belonging to FLA subgroup B (FLA15-FLA18) had reduced levels in *galt7galt8* mutants. The function of FLAs in plants remains largely unknown. The fasciclin-domain is an evolutionarily conserved structural motif originally characterised as a cell adhesion domain in animals (Jay and Keshishian, 1990; Seifert, 2018). The Arabidopsis B-group FLAs are characterized by the lack of a GPI anchor sequence and two tandem fasciclin domains which are separated by a proline, alanine, serine and threonine (PAST)-rich region (Johnson et al., 2003). The lack of a GPI anchor implies that the B-group FLAs are secreted into the apoplasmic space (Showalter, 2001), but a recent study indicated that FLA16 is located both at the plasma membrane and the cell wall (Liu et al., 2020). The location of FLA16 at plasma membrane-cell wall interface suggests that it could interact with proteins involved in cellulose biosynthesis. The PAST-rich domain is thought to contain AGP-like glycans since such epitopes were identified on Arabidopsis FLA4-citrin fusion proteins when using AGP glycan-specific antibodies LM14 and JIM13 (Xue et al., 2017). Mutations in the predicted AGP motifs from FLA4-citrin altered the post-secretory fate of the protein and resulted in a reduced FLA4-citrin signal at the plasma membrane. Similar fates can be envisioned for FLA15-18 in the *galt7galt8* mutants, and could explain the defective cellulose biosynthesis observed for these mutants. It should be noted that the reduction in cellulose biosynthesis is unlikely triggered by a signalling cascade that regulates *CESA* transcription, since the *galt7galt8* mutants did not show any significant changes in either the primary or secondary cell wall *CESA* transcript levels.

It is possible that GALT7 and GALT8 can synthesize galactans by directly attaching glycans to CESAs or other proteins involved in cellulose biosynthesis. The possibility of CESA *N*-glycosylation has been investigated before, but considered unlikely since no molecular weight differences were observed between the CESAs from the PNGase-treated protein extract of the *N*-glycosylation deficient mutant *cgl1* and the CESAs from untreated WT plants (Gillmor et al., 2002). In the present study, we did not observe molecular weight shifts in the polyacrylamide gel (PAGE) electrophoresis of secondary cell wall CESAs that would support differential glycosylation in the *galt7galt8* background (Supplementary Fig. 8). Of the proteins known to participate in cellulose biosynthesis, the plasma membrane-associated, GPI-anchored COBRA (COB) protein and KOR1 have been shown to be *N*-glycosylated (Roudier et al., 2005; von Schaewen et al., 2015). COB and KOR1 do not contain AGP-like *O-*glycosylation motifs, but their AG glycosylation cannot be excluded. AGP *O-*glycosylation motifs are not fully understood, as shown by the presence of AGP glycans on the mutant form of FLA4 which lacks all of the predicted AGP glycosylation motifs (Xue et al., 2017). Interestingly, a reduction in cellulose biosynthesis was also observed in mutants lacking the Golgi-localized STELLO proteins from the GT75 family (Zhang et al., 2016). STELLO proteins interacted with CESAs in yeast two-hybrid experiments and bimolecular fluorescence complementation assays in tobacco leaf epidermis cells (Zhang et al., 2016). The Golgi distribution, secretion and plasma membrane velocity of the CSC was impaired in the *stello* null mutant, which translated to reduced cellulose biosynthesis rates. The GT75 domain was shown to be essential for STELLO function, but the enzymatic activity of these proteins has not yet been established. It is hypothesized that STELLO may glycosylate CSCs either directly or indirectly by targeting components that affect CSC function and localization (Zhang et al., 2016).

Based on our results, we hypothesise that the reduced level of FLA subgroup B observed in *galt7galt8* mutant is linked to reduced CESA protein levels, which leads to decreased cellulose biosynthesis rates and the further decrease in total β-Yariv accessible AGPs. Orthologs of Arabidopsis FLA7 and FLA11 have been found to directly interact with secondary cell wall CSCs in cotton fibers and the differentiating xylem of *Populus deltoides x trichocarpa* based on co-immunoprecipitation results with secondary cell wall CESA-specific antibodies (Song et al., 2010; Li et al., 2016). We speculate that a GALT7/8 – FLA subgroup B pathway affects how FLA subgroup B members interact with the CSC – or the associated proteins – which, in turn, affects CESA levels and cellulose biosynthesis. Alternatively, or in addition, the FLA subgroup B may facilitate cellulose microfibril assembly and deposition in the cell wall as the cellulose microfibril emerges from the CSC. The recent study of a FLA subgroup B (*fla16*) mutant with a 79% reduction in the expression of *FLA16* revealed a similar growth phenotype as what was witnessed in the case of *galt7galt8*, i.e., reduced inflorescence stem length, shorter etiolated hypocotyls and decreased cellulose biosynthesis in mature stems (Liu et al., 2020). However, the growth defects are more pronounced in the *galt7galt8* mutants, a result which could be explained by the fact that this mutant affects all the members of FLA subgroup B.

Interestingly, both GALT7 and GALT8 contain a domain of unknown function (DUF4094) in their N-terminus (Supplementary Fig. 10). The GT31 family enzymes from *Homo sapiens* contain a N-terminal galectin domain involved in substrate recognition (Barondes et al., 1994; Dodd and Drickamer, 2001). The galectin domain is an evolutionarily conserved carbohydrate recognition domain that can bind β-galactosides. However, in Arabidopsis Clade I-III GT31s, the galectin domain has been replaced by DUF4094 (Qu et al., 2008). We speculate that this domain could be involved in recognizing targets or specific glycosylation structures, which would be pivotal to the galactosyltransferase activity of GALT7/GALT8 on specific targets. This type of mechanism could explain the reduction of the entire FLA subgroup B levels in the *galt7galt8* mutants. However, further experimental work will be necessary to establish whether FLA subgroup B and/or other proteins are direct targets of GALT7/GALT8 activity *in vivo*.

## Methods

### Plant material and growth conditions

The Arabidopsis (*Arabidopsis thaliana*) ecotype Columbia (Col-0) was used as a wild-type control. The following Arabidopsis T-DNA insertion lines (Col-0) were ordered from the Nottingham Arabidopsis Stock Centre: *galt8-1* (SALK_110901); *galt8-2* (SALK_150336); *galt7-1* (SALK_069854); *galt7-2* (SALK_053895); CESA4 null mutant (SALK_084627; *irx5-4*); CESA7 null mutant (GABI_819B03; *irx3-7*); and CESA8 null mutant (GABI_339E12; *irx1-7*). The T-DNA location of the *GALT* lines was determined by polymerase chain reaction (PCR) and sequencing using gene- and T-DNA specific primers (Supplementary Table 3). Plants were grown on soil at 22 °C with a photoperiod of 16 h light and 8 h dark at 65% relative humidity. The plants were exposed to a light intensity of 150 µmol m^-2^s^-1^, which was emitted by Valoya NS12 light emitting diode (LED) tubes (Valoya Oy, Helsinki, Finland).

### Plant vector construction and transformation

The genomic sequences of *GALT7-YFP* and *GALT8-YFP* were amplified using primers listed in Supplementary Table 3. The amplified DNA fragments were cloned into the pDONR207 plasmid via Gibson assembly and recombined into pHGY (Kubo et al., 2005). Stable transgenic *galt7-1galt8-1* plants carrying either pGALT7:*GALT7-YFP* or pGALT8:*GALT8-YFP* were generated by *Agrobacterium tumefaciens*–mediated transformation, described in more detail by Nibbering et al. (2020). Transgenic seedlings were selected by hypocotyl elongation on ½ Murashige and Skoog (MS) plates supplemented with 30 μg/mL hygromycin.

### Hypocotyl elongation experiments

Seeds were surface-sterilized with 70% ethanol and stratified for three days on ½ MS agar plates (0.9% agar, 5 mM MES, pH 5.7). The seeds were exposed to light for six hours and then placed in the dark for four days. The plates were scanned and hypocotyl elongation was quantified with Image J.

### Light microscopy and Transmission electron microscopy (TEM)

For the light microscopy, the bottom 1 cm of 10-week-old inflorescence stems was cut into 70µm cross sections with a vibratome. The cross sections were stained for 1 minute in 0.01% toluidine blue and washed twice with water. Images of the cross sections were captured using an axioplan microscope with a camera attachment.

For the TEM, the bottom 1 cm of 10-week-old inflorescence stems was cut into small pieces and fixed in 2.5% Glutaraldehyde (TAAB Laboratories, Aldermaston, England), 4% paraformaldehyde in 0.1M sodium cacodylate buffer, and further postfixed in 1% aqueous osmium tetroxide (TAAB Laboratories). Samples were dehydrated in ethanol and propylene oxide and finally embedded in Spurr’s resin (TAAB Laboratories). All of the samples were processed using a Pelco Biowave Pro+ microwave tissue processor (Ted Pella, Redding, CA). The 70nm ultrathin sections were post-contrasted in uranyl acetate and Reynolds’ lead citrate and further examined with a Talos L120C (FEI, Eindhoven, The Netherlands) operating at 120kV. Micrographs were acquired with a Ceta 16M CCD camera (FEI, Eindhoven, The Netherlands) using TEM Image & Analysis software (version 4.17; FEI, Eindhoven, The Netherlands).

### Wet chemical analysis of cell walls

The bottom 10 cm of 10-week-old soil-grown plants was lyophilized and ground with a ball mill. The alcohol insoluble residue (AIR1) and starch (AIR2) were removed according to a protocol described by Nibbering et al. (2020). Crystalline cellulose was quantified according to a methodology presented by Updegraff (1969). The monosaccharide composition was determined according to the methodology of Latha Gandla et al. (2015), while lignin content was quantified with a lignin standard according to the protocol described by Fukushima and Hatfield (2004).

### AGP analysis using β-Yariv precipitation

β-yariv was synthesized according to the methodology reported by Yariv et al. (1962). The first 10 cm of inflorescence stems, four-day-old hypocotyls and seven-day-old seedlings was harvested, lyophilized and ball milled. The amount of AGPs in 1.5 mg of dry weight sample was quantified according to the protocol described by Lamport (2013) via a gum Arabic standard. The β-yariv gel diffusion and electrophoresis was performed according to the protocol described by Veenhof and Popper (2020). Briefly, soluble AGPs were extracted by adding 50 µl demineralized water per mg of dry weight. The samples were incubated for 30 min at 50 °C and vortexed every 10 minutes. The samples were spun down at full speed for 2 min and the supernatant was transferred to a new tube. The supernatants were concentrated using a SpeedVac until a desired volume was reached. For the β-yariv gel diffusion, soluble AGPs from a 3 mg dry weight sample were loaded to the gel. For the β-yariv gel electrophoresis, soluble AGPs from a 10 mg dry weight sample were loaded to the gel.

### Visualisation of co-localization using confocal microscopy

The lines *pGALT7:GALT7-YFP* and *pGALT8:GALT8-YFP* co-expressing the marker lines SYP32-mCherry or SYP43-mCherry were generated by plant crossing, and imaged according to the methodology presented by Nibbering et al. (2020).

### *CESA* expression and CESA Western blot analysis

Four-day-old dark-grown hypocotyls and the first 10 cm of 20 cm inflorescent stems were flash-frozen in liquid nitrogen and ground with a mortar and pestle. Total RNA was extracted according to the protocol of the TRIzol RNA Isolation Reagent(Simms et al., 1993). Any potentially contaminating DNA was removed with DNase treatment, after which cDNA was synthesized using the Maxima First Strand cDNA Synthesis Kit (Thermo Scientific, Waltham, MA). CESA expression was measured using *CESA* and *UBQ1* specific primers according to the protocol of Wang et al. (2015).

Proteins were extracted from a 30 mg fresh weight sample based on the methodology presented by Hill et al. (2014). A total of 10 µg of protein was loaded into every well of the SDS-PAGE gel. Following electrophoresis, the proteins were transferred to a PVDF membrane. The membranes were blocked in 5% milk powder in TBST. The membranes were blotted with the primary antibodies CESA4 (Agrisera, AS12 2582), CESA7 (Agrisera, AS12 2581) or CESA8 (Agrisera, AS12 2580), prepared as 1:1000 dilutions in 2.5% milk powder in TBST. The membranes were then blotted with the goat anti-rabbit IgG antibody (H+L), peroxidase PI-1000 (vector), prepared as a 1:10000 dilution in 2.5% milk powder in TBST.

### ^13^C labelling of Arabidopsis seedlings

Seeds (20 mg) were surface-sterilized for 5 min in 70% ethanol, 10 min in chlorine solution, and then washed three times for 5 min in sterile demineralized water and stratified for three days at 4°C. The seedlings were grown for eight days in 50 ml ½ MS media (5 mM MES, 1% glucose, pH 5.7) in 250 ml flasks with a photoperiod of 16h light (24°C) and 8 hours dark (18°C). On the eighth day, 1% glucose (90% glucose, 10% ^13^C-glucose, Sigma Cas number: 201136-45-0) was added to the seedlings and samples were taken 0, 8, 16 and 24 hours after addition of glucose. The seedlings were washed twice in demineralized water and flash-frozen in liquid nitrogen. The seedlings were lyophilized and ground with a ball mill (total dry weight). The AIR1/AIR2 fractions were removed from the dry cell wall powder and the crystalline cellulose was extracted according to the protocol presented by Updegraff (1969). From the total dry weight and crystalline cellulose, 5 mg was weighed down, packed and measured with an elemental analyzer (Flash EA 2000, Thermo Fisher Scientific, Bremen, Germany) coupled to an isotope ratio mass spectrometer (DeltaV, Thermo Fisher Scientific, Bremen, Germany) to determine the ^13^C content (Werner et al., 1999). The ^13^C fraction of total carbon was calculated from working standards of wheat and maize flours calibrated against IAEA-600, IAEA-CH-6, and USGS40 reference standards.

### Global Proteomics

The first 10 cm of 20 cm inflorescent stem was flash-frozen in liquid nitrogen and ground with a mortar and pestle. Proteins were extracted and trypsin-digested following procedures described by by Zhang et al. (2018), followed by a post digestion clean-up on an Oasis® HLB µElution plate (Waters, Massachusetts, USA). Approximately 0,5 µg of each sample were loaded on a BEH C-18 analytical column (75 μm i.d. × 250 mm, 1.7 μm particles; Waters, Massachusetts, USA), and separated using a concave 120 min gradient of 1-40 % solvent B (0.1% formic acid in acetonitrile) solvent A (0.1 % aqueous formic acid) at a flow rate of 300 nL min^-1^. The eluate was passed to a nano-ESI equipped SynaptTM G2-Si HDMS mass spectrometer (Waters, Massachusetts, USA) operating in sensitive mode. All data were collected using ion-mobility MSE with dynamic range extension enabled using a scan-time of 0.4 sec, mass-corrected using Glu-fibrinopeptide B as reference peptide. The data were processed with Progenesis QI software used for quantification (Waters, Milford, MA), and the resulting spectra were searched against the database Araport11 (The Arabidopsis Information Resource TAIR).

### Targeted CESA proteomics

Unique peptides were selected as quantitative surrogates for CESA4 (AT5G44030, APEFYFSEK), CESA7 (AT5G17420, ATDDDDFGELYAFK) and CESA8 (AT4G18780, DIEGNELPR) and analysed as described in Zhang et al. (2018) An isotope labelled peptide for CESA4 (FDGID^**13**^**L**ANDR) was also used in the analysis. Samples were processed as for global proteomics described above. For LC-MRM, samples were analyzed by reversed-phase liquid chromatography-electrospray ionization-MS/MS using a nanoACQUITYTM UPLC (Waters, USA) system connected to a triple-quadrupole mass spectrometer (Xevo TQ-S, Waters, USA) in trapping mode. The trapping flow rate was 5 µL/min and trapping time was 3 min. The separation cycle time took 32 min at a flow rate of 2 µL/min. Peptides were loaded onto BEH iKey (C18, 1.8 µm, 150 µm i.d. × 50 mm; Waters) in 99% solvent A (0.1% FA in H_2_O) and 1% of solvent B (0.1% FA in acetonitrile). The percentage of solvent B was elevated from 1% to 30% in 18 min linear gradient, then increased from 30% to 40% solvent B in 2 minutes, followed with 2.5 min linear gradient to 95% of B and held at 95% for 2.5 min. After that, the column was balanced by ramping to 1% solvent B in 0.5 min and flushing for 7 min. Data acquisition was performed using the following parameters: capillary voltage, 3.2 KV; source temperature, 120°C; cone gas, 60 L/h. 5 transitions per peptide were monitored in a scheduled way, with a cycle time of 2 sec and a 4 min MRM detection window. The cone voltage and collision energy for each transition were done according to by Zhang et al. (2018). Data were imported into Skyline 4.1.0 (MacCoss lab software, Washington University, USA) with Savitzky-Golay smoothing. All peaks were manually inspected to ensure correct peak detection and accurate integration.

## Author contributions

P.N., R.C., G.W. performed experiments and analyzed data. P.N. and T.N. planned experiments and analyzed data. P.N. and T.N. wrote the manuscript with help from the other authors.

## Acknowledgements

We would like to thank Dr. Junko Takahashi-Schmidt and the UPSC Biopolymer Analytical Platform for help with the cell wall monosaccharide analysis. We would like to acknowledge the facilities and technical assistance of the Umeå Core Facility Electron Microscopy (UCEM) at the Chemical Biological Centre (KBC), Umeå University, a part of the National Microscopy Infrastructure NMI (VR-RFI 2016-00968). This work was supported by The Swedish Foundation for Strategic Research (Value Tree), Bio4Energy (Swedish Programme for Renewable Energy), the UPSC Centre for Forest Biotechnology funded by VINNOVA and the Swedish Research Council for Sustainable Development (Formas).

